# Clay-induced DNA double-strand breaks underlay genetic diversity, antibiotic resistance and could be a molecular basis for asbestos-induced cancer

**DOI:** 10.1101/111534

**Authors:** Enrique González-Tortuero, Jerónimo Rodríguez-Beltran, Renate Radek, Jesús Blázquez, Alexandro Rodríguez-Rojas

## Abstract

Some natural clays and synthetic nanofibres present in the environment have a severe impact on human health. After several decades of research, the molecular mechanism of how asbestos induce cancers is not well understood. Different fibres, including asbestos, can penetrate the membrane and introduce DNA in both, bacterial and eukaryotic cells. Incubating Escherichia coli with sepiolite, a clayey material, and asbestos under friction forces, both fibres cause double-strand breaks in bacteria. Since antibiotics and clays are used together in animal husbandry, the mutagenic effect of these fibres might constitute a pathway to antibiotic resistance due to the friction provided by peristalsis of the gut from farm animals in addition to the previously proposed horizontal gene transfer. Moreover, we raise the possibility that the same mechanism could generate bacteria diversity in natural scenarios with a role in the evolution of species. Finally, we provide a new model on how asbestos may promote mutagenesis and cancer based on the observed mechanical genotoxicity.

## Introduction

Clays such as sepiolite are jointly used with antibiotics in farming as growth promoters. This practice improves growth and animal product quality, and these additives are common in feed for broiler chickens and pigs [1,2]. Sepiolite is considered to be safe, stable and chemical inert hence being also used in tablet formulation for human medicine [3]. However, in a recent study clays used as animal feed additive can increase the risk of horizontal gene transfer (HGT) among microbes, resulting possibly in a rise of antibiotic resistance [4,5].

In this case, the transformation of bacteria by foreign DNA can be achieved when clay fibres are spread by friction or vibrations. This phenomenon is known as Yoshida effect [6] and relies on the ability of mineral nanofibres or nano-needles to adsorb DNA and to penetrate bacterial cells under sliding friction forces [7]. By its mechanical nature, the Yoshida effect can be used to transform diverse bacterial species [5,8,9]. The action of sepiolite and other clays fibres is not only capable of delivering DNA into the receptor bacteria but also able to promote the releasing of DNA by disrupting the cell envelope of the portion of the population by the abrasive action of clays [5].

Before Yoshida began his experiments with bacteria, the ability of asbestos to transform eukaryotic cells was reported at the end of the eighties [10]. In fact, fibrous clays and industrial nanofibres are considered genotoxic and carcinogenic, likely due to their ability to damage DNA [11]. They have assayed in several experimental models including bacteria and cell in cultures, but they display a poor correlation with mutagenicity or carcinogenesis found *in vivo* [12,13]. According to these observations, a significant concern arises from fibrous clays or industrial nanofibres which are responsible for severe human diseases such as asbestosis [14]. However, short or long periods of exposure to fibres have been failing to identify a molecular basis of DNA damage in different several genotoxicity tests [14]. Thus, nowadays the mechanisms underlying the genotoxicity and carcinogenicity of asbestos and other fibres remain obscure.

Additionally, clays may have the potential to enhance antibiotic resistance in farming activities [4]. In natural scenarios, sediments and stones (gastroliths) are frequently swallowed by animals resulting unavoidably in the exposure of their microbiota to pebbles, sand, and clays. Soils and waters are a primary source of antimicrobials, either by natural microbial production or environmental antibiotic pollution, a major selective pressure that favours resistant strains [15,16]. Even, gut microbes can produce antibiotic compounds [17].

In this study, the ability of fibrous clays such as sepiolite and asbestos to transform bacteria and to induce mutagenic DNA double-strand breaks (DSBs) when they are exposed to friction forces was experimentally shown. Additionally, a molecular mechanism of action for asbestos, which was a strong inducer of DSBs in *Escherichia coli* when friction is present, was proposed. Finally, the importance of this mechanism is discussed for the speciation processes of animals that use gastroliths.

## Results and Discussion

### Mutant frequency in sepiolite-treated cells is higher than in non sepiolite-treated cells

Different types of clays can transform bacteria by absorbing DNA and penetrating the cell envelope. In that case, the penetration could allow the clays to interact with the intracellular DNA and promote mutations. To test whether sepiolite under friction forces (as in transformation) has an impact on bacterial mutation rate, the mutant frequency of *Escherichia coli* was measured by plating in the antibiotic fosfomycin and enumerating spontaneous mutants (fig 1). When the cells were merely exposed to sepiolite without any friction on agar plates surface, no significant differences in mutant frequencies were detected (Mann-Whitney U test; P=0.999). In contrast, a six-fold increase in mutant frequency was found when friction was present for two or three minutes (Mann-Whitney U test; P=0.008) and a modest increase—but not significant—when the treatment time lasted for one minute (Mann-Whitney U test; P=0.421). Interestingly, only cells in the stationary phase displayed an increase in mutant frequency (Kruskal-Wallis test; P=0.001), while no significant mutagenesis was found when bacteria came from exponential cultures (Kruskal-Wallis test; P=0.954; fig 1). Along with the mutant frequency experiments, the effect of the treatments on cell viability was checked. Sensitivity to the treatment is higher in exponential phase than in stationary phase cultures (fig 2). The observed mutant frequency differences was initially attributed to a higher sensitivity to the treatment. However, several hypotheses can explain the observed increase in mutant frequency and reduced-sensitivity in stationary phase cultures.

**Fig 1.**
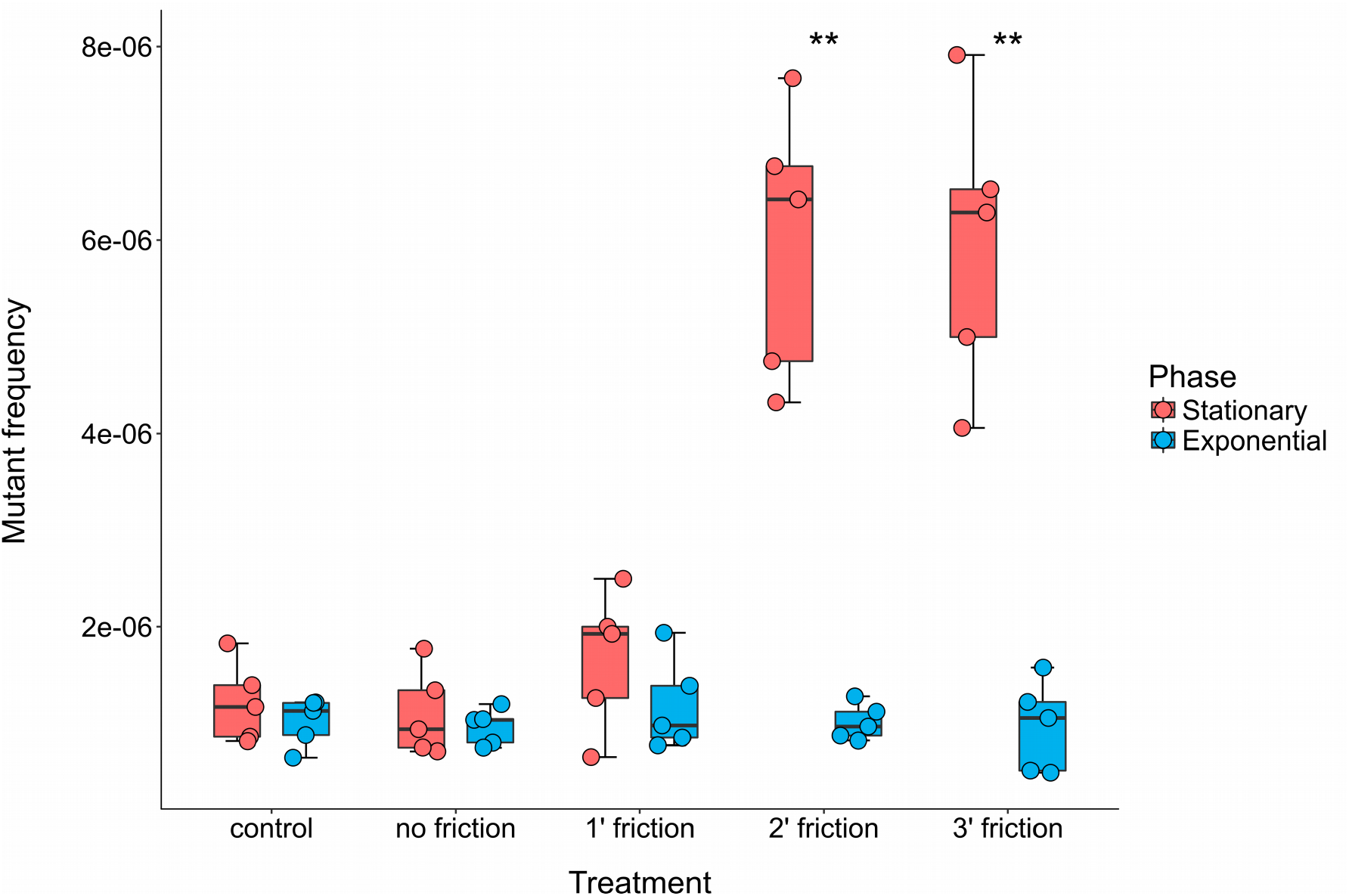
Sepiolite can be mutagenic after friction treatment only in stationary phase. Box-plot of the mutant frequency induced by sepiolite treatment in *E. coli* MG1655 (A) stationary and (B) exponential phase cells. The x-axis indicates the experimental treatment (control, mixture of bacteria and sepiolite without friction force, and with friction force during one, two and three minutes). Asterisks represents significant difference; Mann-Whitney *U*: P < 0.01.

**Fig 2.**
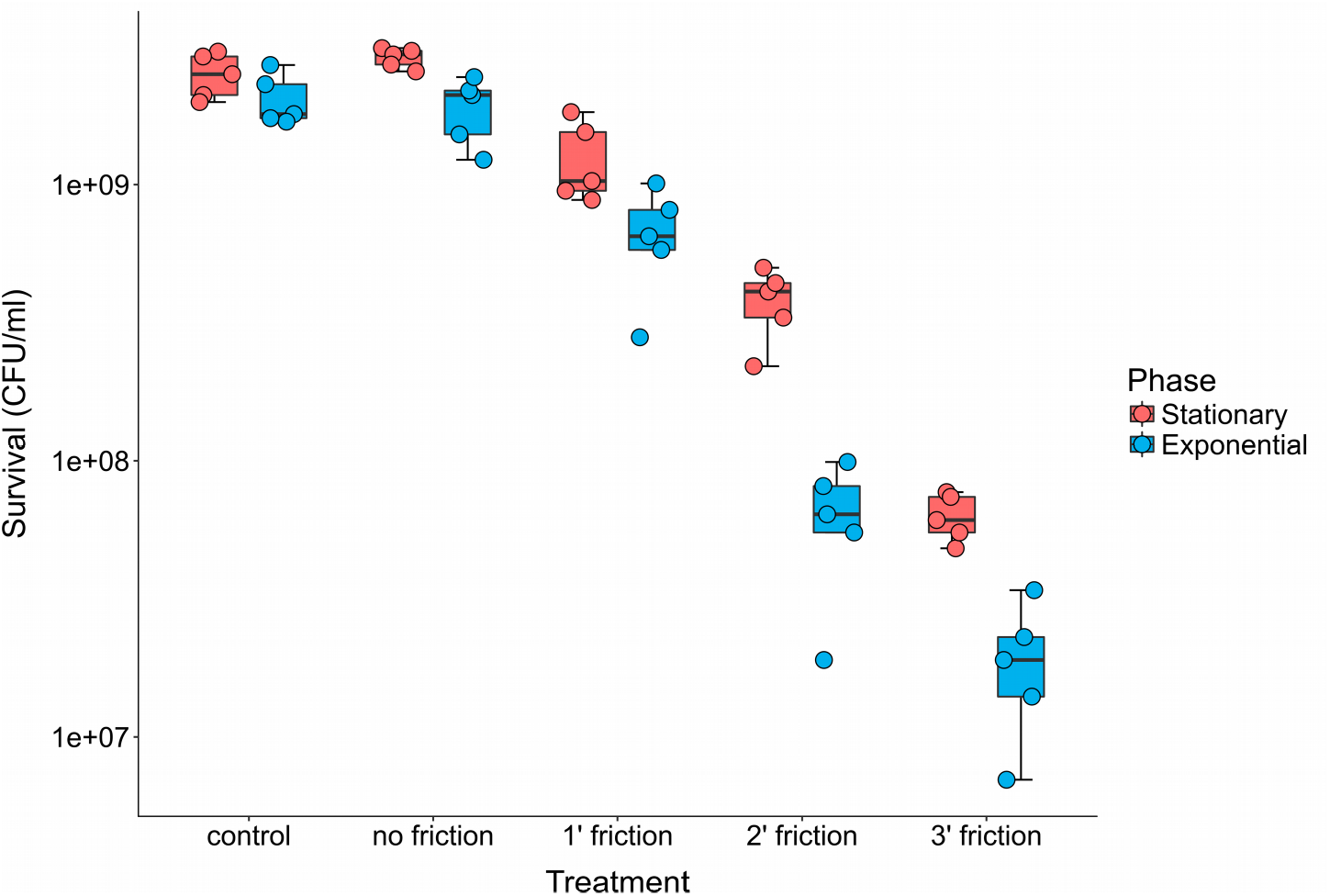
The higher the time of friction, the smaller the cell viability. Box-plot of the survival of *E. coli* MG1655 to the action of friction with sepiolite during one, two and three minutes of treatment. Groups with and without sepiolite gently spread with glass beads onto agar plates were used as controls.

### Heavy metals are non relevant on sepiolite mutagenesis

Many minerals containing metals such as iron, aluminium or copper are toxic for bacteria because of the generation of reactive oxygen species (ROS) via the Fenton reaction [18]. The release of metal ions inside the cell could therefore be the reason of the increase in mutagenesis. In fact, despite the addition of 2-2′ bipyridyl, a chelating agent, shortly before treatment, mutagenesis was still observed (Kruskal-Wallis test; P=0.008; fig S1). This result indicates that the mutagenic effect does not depend on the metals present in the fibres.

### Sepiolite interacts with the DNA by causing double-strand breaks

A second likely explanation is the physical interaction of individual clay fibres in motion directly damaging DNA by creating DSBs. The ability of sepiolite fibres to penetrate and interact with DNA has already been stated [6,19]. Physical or mechanical stress on the DNA duplex is a relevant cause of DSBs [20]. To evaluate this possibility, *E. coli* DH5α strain (*recA* deficient) carrying the plasmid (pET-19b) was subjected to treatment with sepiolite and sliding friction. Sepiolite without friction and bacterial cells alone were used as controls. The plasmid content was extracted, and its integrity was evaluated by gel migration (fig 3A). Typically, during plasmid DNA extraction, three molecular conformations are found: the supercoiled (which migrates very fast), nicked DNA (which is also closed circular but relaxed due to single strand breaks and it has an intermediate migration rate) and linear molecules (with a lower migration speed) [21]. These latter DNA molecules were especially abundant in the friction-sepiolite treated group at the time that they are present in a low level in control groups. In fact, plasmids from the sepiolite group (under friction) presented a significantly high level of linearised molecules when compared to the control groups (One-Way ANOVA test; P=3.33×10^−16^). According to these results, the joint action of sepiolite and friction are responsible for induction of DSBs in the DNA. Interestingly, no increase in nicked DNA (single-strand break) was observed, indicating that if this type of lesion occurs, it happens at a non-detectable rate by this technique (fig 3B).

**Fig 3.**
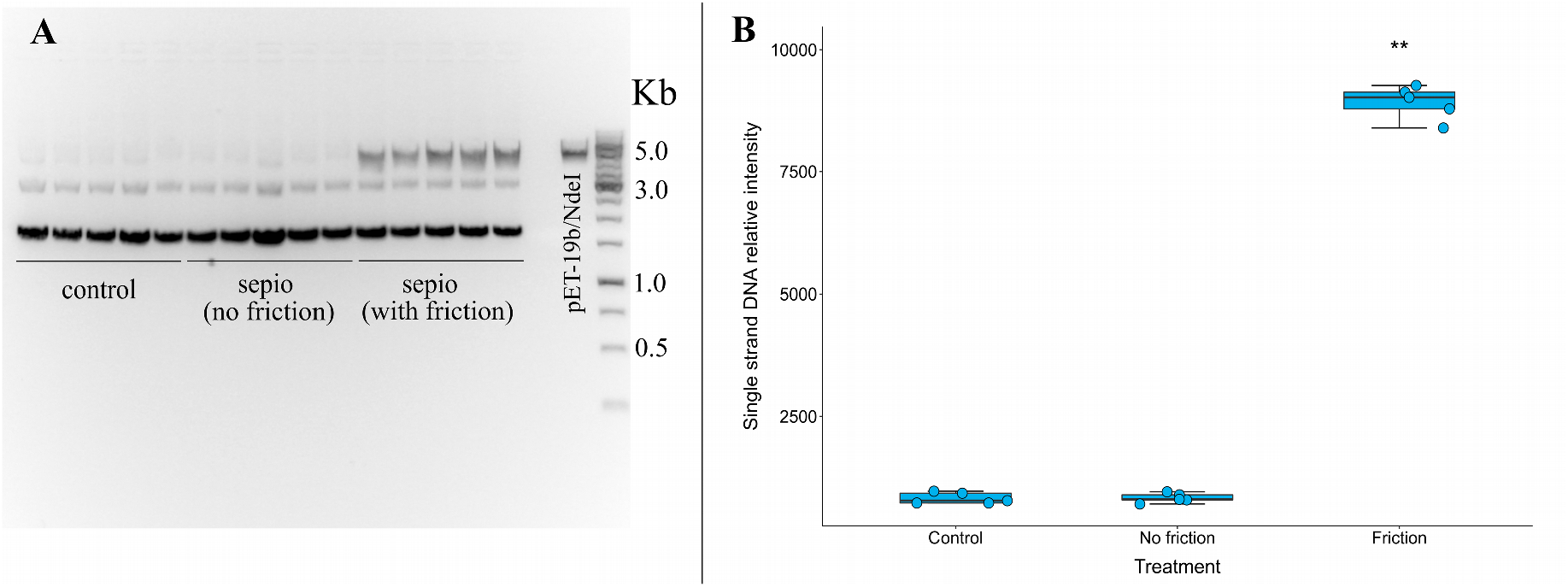
Linear plasmid DNA molecules abundance is higher when friction forces are applied. (A) Extraction of the plasmid pET-19b from sepiolite-treated *E. coli* DH5α, a *recA* deficient strain, during one minute (five extractions per treatment). Note the enrichment in linearised plasmid DNA molecules from bacteria treated with sepiolite under two minutes of friction applied in 1% agarose gel. (B) Box-plot of the abundance of single strand DNA molecules under different experimental treatments (control, sepiolite without friction and sepiolite with friction). Plasmid pET-19b digested with a single cut site enzyme NdeI was used as a control for the linear molecule migration rate and as a reference to calculate relative intensities using a densitometry analysis. Asterisks represents significant difference; Tukey HSD test: P < 0.01.

The view of mutagenic DSBs by mechanical shearing is very consistent with the absence of mutagenic effect in exponentially growing bacteria. If the organism is diploid (even if the diploidy is only transient, as in replicating bacteria or replicating haploid yeast), then homology-directed repair can be used [20]. Because *E. coli* lacks a pathway to join non-homologous ends, homologous recombination is the only mechanism to salvage broken chromosomes [22]. But how can *E. coli* repair DSBs in stationary phase by homologous recombination? Stationary-phase cultures contain cells with several chromosome copies [23]. In exponentially growing *E. coli* DSB repair is non-mutagenic [24,25]. However, break repair becomes mutagenic during the stationary phase and requires the Sigma S factor (RpoS), the SOS response, and the error-prone DNA polymerase PolIV. The change from one situation to the other has been described as a switch from high-fidelity repair in the exponential phase to error-prone DNA double-strand breaks during the stationary phase [24,25]. Because DSBs are lethal unless repaired, and repair action requires RecA protein [24,25], the experiment of sepiolite mutagenesis was repeated with *E. coli* DH5α that is impaired in the SOS response triggering to confirm this notion. In such analysis, sepiolite mutagenesis was completely abolished by *recA* gene inactivation in stationary phase (Kruskal-Wallis test; P=0.011; fig 4). Thus, the lower level of mutant frequency in the *recA* deficient strain could be explained by the death of cells that suffered DSBs and were unable to repair them. Mutations introduced by DSB repair are considered a mechanism of diversity via mutagenic repair in bacteria [26,27].

**Fig 4.**
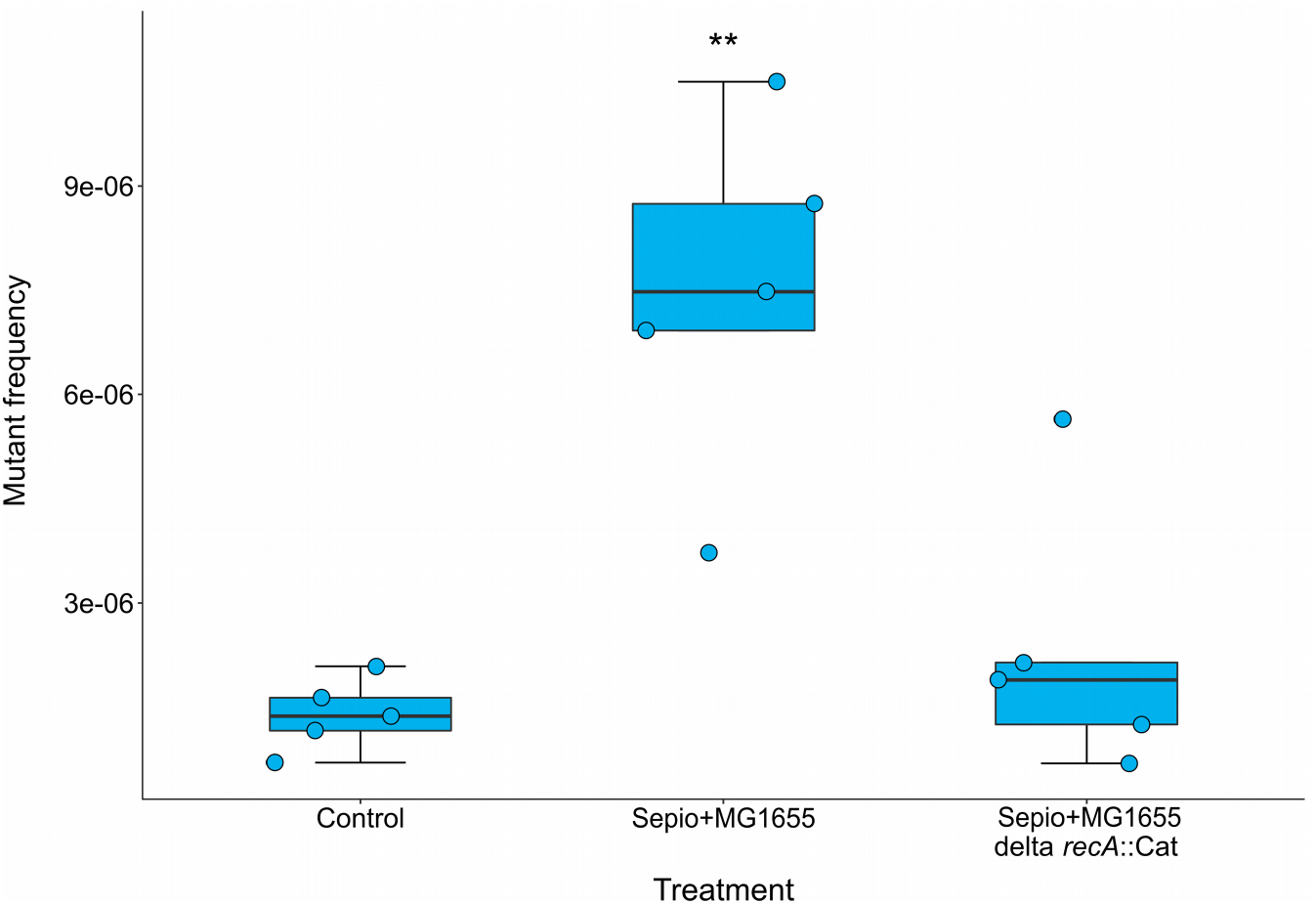
Inactivation of the *recA* gene suppresses the mutagenic effect of sepiolite under friction in *E. coli* MG1655. Box-plot of the mutant frequency of *E. coli* MG1655 and DH5α (derivative *recA* mutant) when treated with sepiolite during two minutes. Asterisks represents significant difference; Mann-Whitney *U*: P < 0.01.

Potentially, the mutagenicity of clay treatment is also enhanced in stationary phase cells due to DNA being more tightly compacted than in the exponential phase [28]. Indeed, in *Escherichia coli*, DNA goes to a co-crystallization state with the stress-induced protein Dps offering protection to several types of stress, ordinarily chemical damage [29]. However, while crystallization is often associated with less flexibility or added fragility to direct physical contact, less compacted DNA of proliferating *E. coli* is elastic and soft [30], which may limit the number of DSBs. It is then possible that mineral fibres under friction can break DNA strands more easily in the stationary than in the exponential phase.

### Sepiolite fibres can penetrate bacteria when friction forces are present

To reunite more evidence that penetration and interaction of fibres with DNA cause DSBs inside the cell, a direct observation of sepiolite-treated bacteria by scanning electron microscopy (SEM) was performed. Fibres look compatible in dimensions able to penetrate bacteria without completely destroying the envelope. Additionally, bacteria were directly penetrated by fibres while those that were exposed to mineral without friction were not (fig 5). This observation is in concordance with previous studies, whereas sepiolite and other nano-sized acicular materials can penetrate bacterial cells under friction forces on a hydrogel [6]. In fact, the partial destruction of the cell wall and the presence of mutants after adding 2-2′ bipyridyl point to the mechanical action as causing agent of the damage. The notion of mechanical breaks is in good agreement with the results in cell-free systems. In these experiments, breakage of plasmid DNA was not directly associated with the amount of iron released by asbestos fibres when they are incubated together [14].

**Fig 5.**
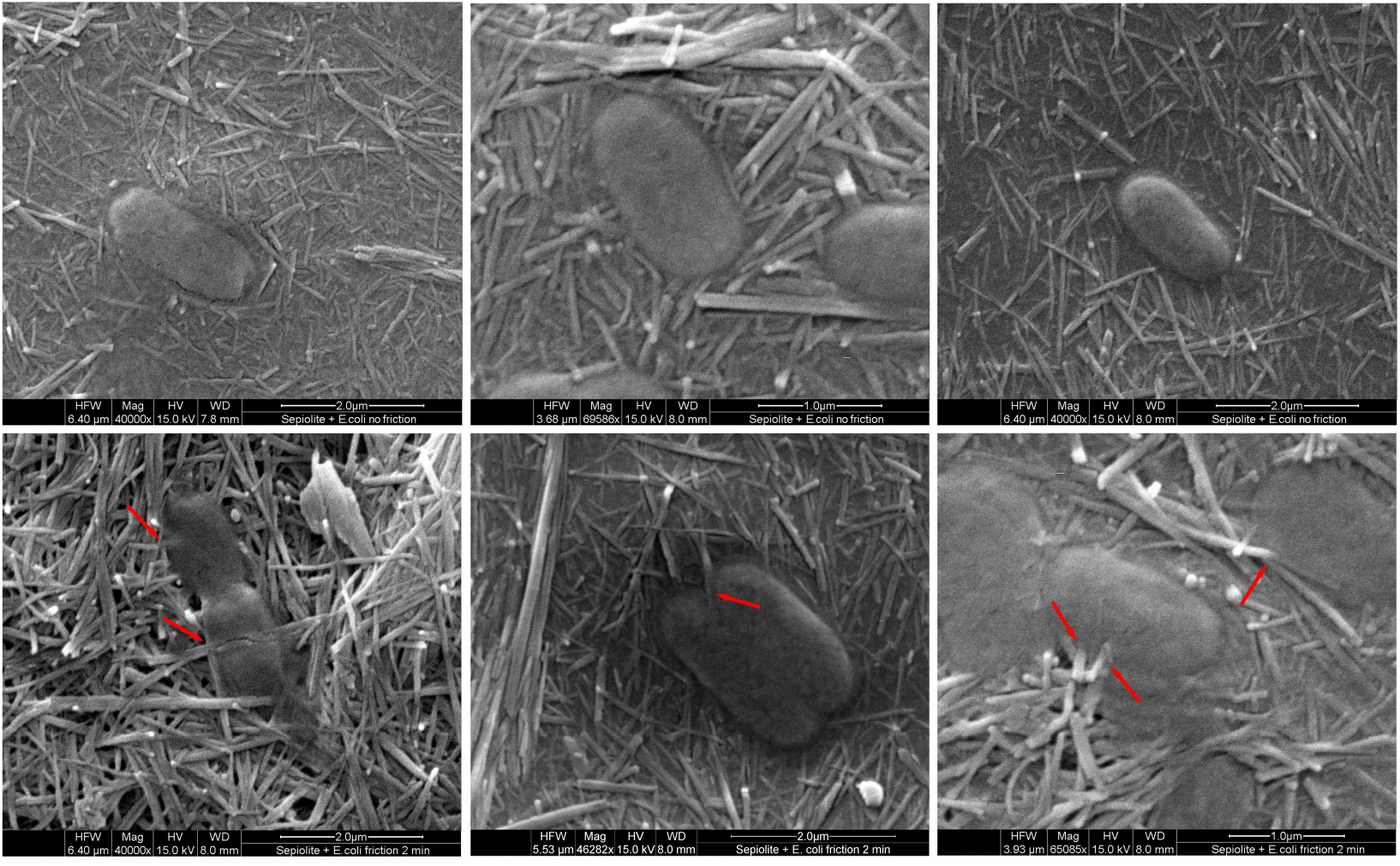
Sepiolite can penetrate bacterial cells when friction forces are applied. SEM of stationary phase *E. coli* MG1655 treated with sepiolite. Red arrows represent potential sites of sepiolite fibre penetration. Bacteria were observed with different magnifications ranging from 40 000X to 70 000X.

### Sepiolite fibre length matters to cause significant DNA damage in the cell

Sepiolite also contains very short fibres (fig S2). In the case of asbestos, there is a certainty that long fibres are much more dangerous by their carcinogenic potential. We designed an experiment to test the influence sepiolite fibre length for mutagenesis in bacteria. The exposure of stationary phase bacteria to a suspension of short fibres (less than 1 μm) did not cause any significant DNA damage when compared with the control and in contrast with the long-fibre original mineral suspension (Kruskal-Wallis test; P=0.005; fig 6).

**Fig 6.**
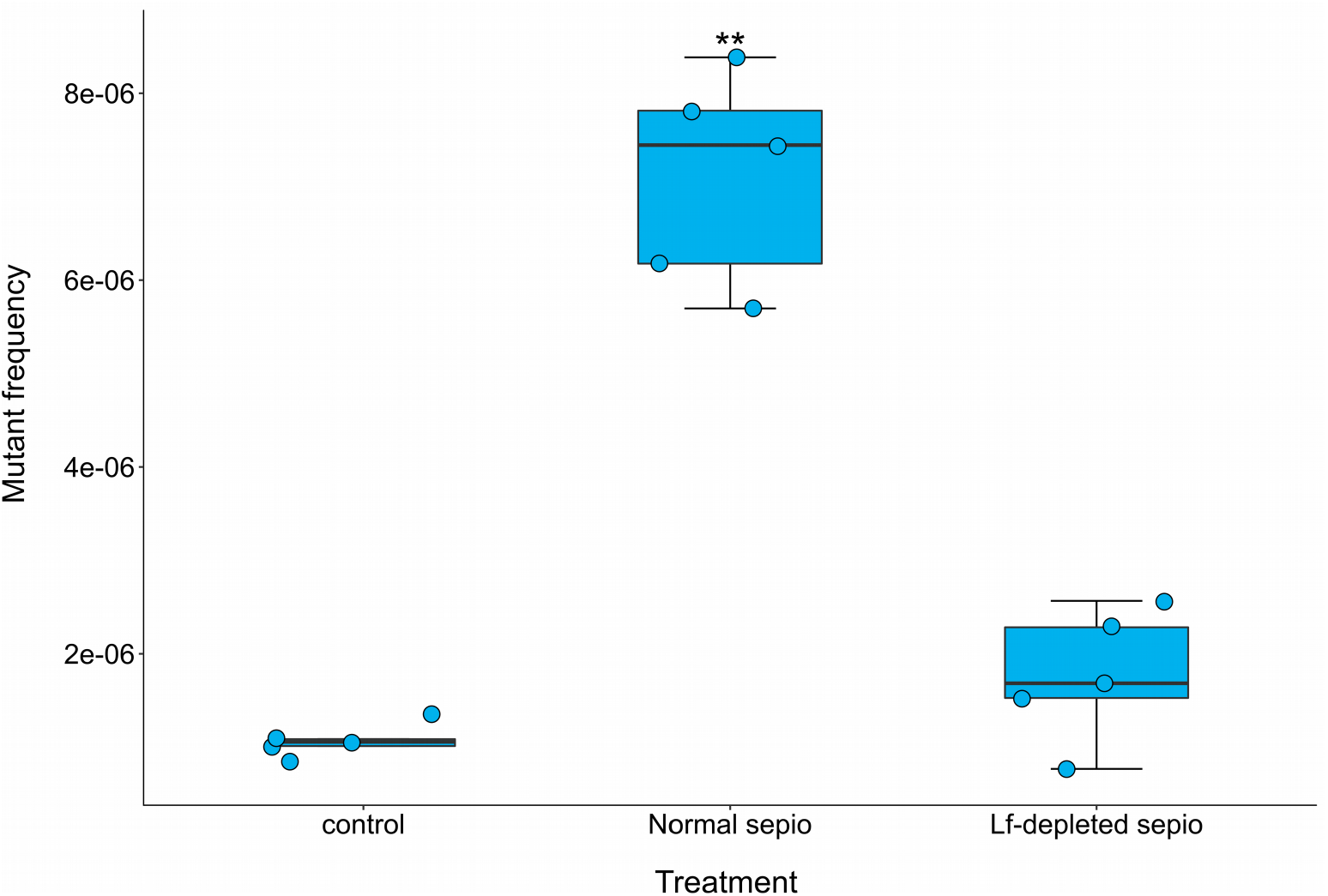
Removal of sepiolite fibres longer than 1 μm decreases fibre-induced mutagenesis to the level of the control. Box-plot of the mutant frequency of *E. coli* MG1655 when sepiolite fibres longer than 1 μm were removed in mutagenesis experiments (lf-depleted sepiolite). Dry and reconstituted sepiolite (normal sepiolite) and bacterial cells (labelled as control) with no sepiolite were used to compare the effects of long fibre removal. Asterisks represents significant difference; Mann-Whitney *U*: P < 0.01.

### Asbestos fibres increase the mutant frequency in the same way as sepiolite do

Bacterial genotoxicity experiments are considered a key step in the assessment of mutagenic properties of chemicals, drugs or materials in general [31]. Because asbestos fibres resemble sepiolite ones, an experiment to test if asbestos fibres provoke an increase in mutagenesis was designed using crocidolite asbestos (fig S2). In our assay, the addition of asbestos to bacteria in the plates without friction did not increase the mutant frequency. In contrast, the application of friction when the fibres were present increased the mutant frequency even more than sepiolite alone (Kruskal-Wallis test; P=0.002; fig 7), probably by the same mode of action. Yoshida *et al*. have suggested that asbestos and other clays can be potentially mutagenic based on integrity analysis of genomic DNA from treated bacteria [32]. A clear antecedent of the ability of fibrous nanoclays to penetrate bacteria was the transformation of monkey cells in culture by exogenous plasmid DNA using chrysotile (a type of asbestos) [10]. Although procedures are not described in details, we think that this transformation requires penetration of the cell membrane.

**Fig 7.**
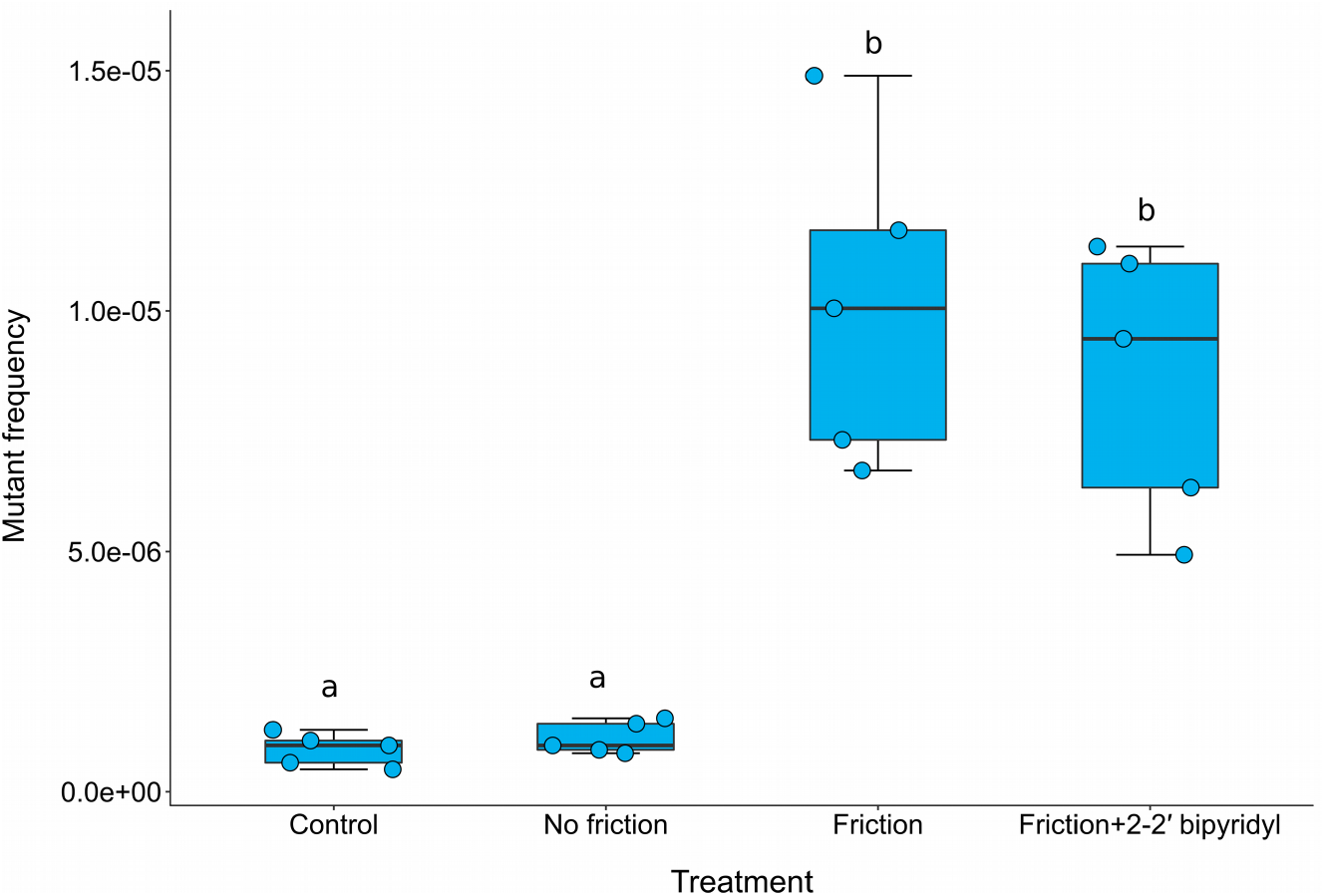
Asbestos can increase the mutant frequency of *E. coli* after friction around one order of magnitude. Box-plot of mutant frequency induced by asbestos (crocidolite fibres) treatment in *E. coli* MG1655. Asterisk represents significant difference; Mann-Whitney *U*: P < 0.05. Equal letters represent no differences while different ones represent significant differences.

### Further discussion

The poor correlation between DNA damage *in vivo* and *in vitro* described in previous studies [12] may be explained by the limited or lack of penetration of asbestos in experimental designs. Thus, the introduction of some friction or shaking can in determining if penetration of cells by asbestos and other fibres underly a molecular mechanism of carcinogenesis. The mechanism(s) underlying asbestos toxicity associated with the pathogenesis of mesothelioma has been a challenge to unravel for more than six decades [33]. According to our results and the current knowledge about asbestos-induced carcinomas, we speculate about a model that explains a potential path leading to carcinomas. Briefly, we think that people exposed to asbestos fibres during prolonged periods accumulate them in the respiratory tract. It is frequent to find asbestos fibres into the pleural cavity, and maybe they increase the friction coefficient in the pleural space, a parameter with a very small value in in physiological conditions [34]. The coelomic movement (a cyclical mechanical movement between the parietal pleura—covering membrane of the inner surface of the thoracic cavity—and the visceral pleura—covering membrane of the lung surface—) provokes the movement of asbestos, trespassing occasionally the mesothelial cell membranes or floating mesothelial cells, physically interacting and disrrupting the DNA or spindle. This physical interaction, with adequate intensity, could induce DSBs, which generate chromosome aberrations or fragmentations in eukaryotic cells as we found here for bacteria. After years of exposure, DSBs or spindle disruption can cause chromosome damages or losses or aneuploidy that increase the probability of malignancy. The proposed model for eukaryote cells would need *in vitro* validation with epithelial cells but this is beyond the scope of the current study and left for future research. Moreover, this model does not exclude other toxic and genotoxic mechanisms of asbestosis such as reactive species arising from metal action or inflammatory response.

One of the most important limitations of our study is the lack of an animal model to test if our finding of mutagenicity in bacteria by clays occurs *in vivo*. In theory, clays present in livestock feed could promote antibiotic resistance and virulence in pathogenic bacteria by not only their transformation ability but also via mutations. However, testing conditions are hindered by the fact that experiments would require at least S1 security level, and this is difficult to achieve with livestock animals [4]. Transformation of plasmid DNA requires penetration and sepiolite and other clays have shown this capacity in a wide range of concentrations although it diminishes at high concentration due to the killing of bacteria [9,35–37]. In a previous study, the values of pressure in the gut of many animal species were discussed, meeting the criteria very well [4]. The presence of a hydrogel does not seem to be a problem since both mucin layer of the gut or mucoid secretion in the respiratory tract can play that role, particularly if fibres have the capacity to change viscosity locally or gradients of viscosity exist across these body compartments.

An implication of our study is the consideration of other factors (such the friction forces) in assessing of genotoxicity and carcinogenesis by certain fibrous materials. Until now, many studies associate clay-induced damage mostly with ROS [14]. DNA damage can be produced by oxido-reduction processes generated by metal containing-fibres. Asbestos fibres are carcinogenic for both, humans and experimental animals, because asbestos produce DNA breaks leading to the formation of micronucleus (a type of chromosomal aberration) [38]. This kind of damage seems to be caused more by mechanical action rather than ROS generation, which can worsen the situation but not necessarily has to be determinant. In other words, we think that ROS is more a symptom than a cause. Another example of a potentially dangerous material are the carbon nanotubes (CNTs), a novel industrial material with many applications. The genetic alterations provoked by these nanotubes in rat malignant mesothelioma were similar to those induced by asbestos [39]. Interestingly, CNTs lack heavy metals in their composition. The nanoscale size and needle-like rigid structure of CNTs appear to be associated with their pathogenicity in mammalian cells [38]. Coincidentally, CNTs can be used to transform bacteria with plasmids [40] in a similar fashion that asbestos [10,41] and sepiolite do [6,42]. It would not be surprising that all these fibrous nanomaterials share their ability to mechanically induce DSBs.

Recently, a possible link between talcum powder and ovarian cancer risk associated with asbestos contamination in talc is under discussion. Although the risk is small, some studies suggested a low or moderate but significant chance of cancer, while other rejected/discarded this correlation [43–45]. It is necessary to advance the understanding of molecular base of DNA damage by asbestos and other industrial fibres. If the proposed model of mechanical/physical DNA breaks is validated in future studies, some genotoxicity assays intended to unveil mutagenic properties of materials (e.g. the test of Ames) should be modified accordingly to include a standardised procedure of friction or promoting some sort of shaking during incubation steps. Similarly, several *in vitro* test, with both bacteria and eukaryotic cells, were modified by researchers and regulatory agencies where introduced the metabolic activation by fraction S9 of liver homogenate [46].

Other implications of the induction of DSBs by nanofibres in bacteria could be related with the microbiota of animals that use gastroliths. It has been suggested that gut microbes play a crucial role in keeping species apart or enhance the speciation [47]. It is tempting to speculate that animals that use gastroliths or sediment ingestion expose their microbiota to the abrasive action of stone derivative fibres. Therefore, the shaping of their own microbes is expected to contribute to their own speciation trajectories. Among animals that use or used gastroliths in their evolutionary trajectories, we find several branches of fishes, amphibians, reptiles (including dinosaurs) and birds. Gastroliths also regularly occur in several groups of invertebrates [48]. Wings (2007) recommends making a distinction between lithophagy and geophagy. Lithophagy (stones larger than 0.063 mm in diameter) is defined as the deliberate consumption of stones that turn into gastroliths after their ingestion. Geophagy is the consumption of soil and it is known for reptiles, birds, and mammals. These soils, rich in clays, salts or fat, serve mainly as a food supplement for the supply of specific minerals or for medical purposes [48]. Both concepts can contribute to getting together all the components that this mechanism needs to operate: gut microbiota, gut mucin mucoid layer (hydrogel) and friction forces provided by the peristaltic pressure of digestive tract in animals, especially the gizzard and the stomach. An interesting question is why sepiolite from limestone gastroliths does not damage the animal gut. A convincing explanation is that the mucoid layer in the gut protects it from the action of these sharp fibres at the time that serve as a protective layer for gut epithelium. In mammals, this mucoid layer is around 200 μm thick and is under continuous renovation [49]. Sepiolite is a natural clay mineral characterised by a nanofibre structure with average dimensions less than or equal to 0.2 micrometers in diameter, and from 2 to 5 micrometers in length, although longer fibres can be present.

### Concluding remarks

Overall, one of the most significant contributions of this article is the proposition for the first time of a bacterial model to test genotoxicity of nanofibres and uncover a new mechanism of action for asbestos that correlates better with *in vivo* observations. Asbestosis is a global health and environmental problem, which molecular basis has been a challenge for several decades [33]. Although asbestos fibres are widely distributed in the anatomy of patients [33,50], the most common cancers caused by asbestos originate in lungs (mostly mesothelioma). If the most explored mechanism of action is based on reactive radicals (chemical damage), why is not there significant differences in the frequencies of other types of carcinoma such as leukaemia, lymphoma, liver or kidney cancer among exposed populations? In the last place, and not less important, is the tighter contact of slippery membranes (a monolayer of flattened epithelial-like cells) of the mesothelium. The pleural space is in continuous movement and constitute preferential target of asbestos-induced carcinogenesis. Of particular interest are free-floating mesothelial cells of the cavity, that even proliferate under damaging conditions [51]. The free-floating cells are the ideal candidates to be penetrated by asbestos in the pleural space. They may be more sensitive to suffer direct (physical) or indirect (chemical) DNA damage and become into a mesothelioma. Finally, sepiolite transformation technique gained some popularity in the last years because there is no need to prepare competence cells [9,19,42,52]. In that case, diverse bacteria can be transformed [4] in both stationary and exponentially growing phases. However, to prevent undesired mutations in both, plasmid and genomic DNA, it is highly recommendable to use exponential phase bacteria, where mutagenesis is not significant, at least in *E. coli*.

## Methods

### Bacteria and growth conditions

The *E. coli* MG1655 wild-type strain and its derivative mutants were cultured in Lysogenic Broth (LB). All experiments were performed at 37°C, with shaking in liquid culture. All solid cultures were grown in LB agar 1.5% for standard procedures and 2% for the sepiolite treatment. All cultures were supplemented with antibiotics when appropriate.

### Mutant frequency estimation of sepiolite treated cells

Approximately 2 × 10^9^ bacterial cells per ml of *E. coli* MG1655 and its derivative mutants from overnight or mid-exponential growing cultures were centrifuged and resuspended in 100 μl of sterilised transformation mixture, consisting of sepiolite (Kremer Pigmente, Germany) suspended in aqueous solution at a final concentration of 0.1 mg/ml. Resuspended cells were spread on plates containing fresh Müller-Hinton-Agar (Sigma-Aldrich, Germany) medium solidified with 2% agar, and Petri dishes were pre-dried in a biological safety flow cabinet for 20 minutes before use. Friction force was provided by streaking bacterial cultures plus sepiolite with sterile glass stir sticks gently pressed onto the medium surface for one, two and three minutes, applying as much pressure as possible without breaking the agar gel. Petri dishes were incubated at 37°C for 2 hours to allow DNA repair if any damage occurred. The plates were gently washed four times with 5 ml of 0.9% sodium chloride solution using a 5 ml pipette. The bacterial suspensions were transferred to 10 ml tubes to recover the cells by centrifugation at 3000 g for 10 minutes. The resulting pellets were resuspended in a final volume of 1 ml of fresh LB an incubate during 1 hour at 37°C to allow the cells to recover. Appropriate dilutions were plated onto LB plates to estimate bacterial viability and in LB plus fosfomycin (50 μg/ml) to estimate the number of resistant mutants. Plates were incubated overnight at 37°C. Each experiment consisted of 5 replicates and was repeated at twice. Mutant frequencies were calculated by using the FALCOR web-tool [53].

### Influence of 2-2′ bipyridyl on sepiolite mutagenesis

The effect of 2-2′ bipyridyl, a metal chelating agent [54], on sepiolite mutagenesis was determined by measuring its influence on the mutant frequency for a selected concentration of sepiolite, where mutagenesis was observed. The experiment consisted of adding a titrating concentration of 2-2′ bipyridyl (200 μM) to chelate metals five minutes before the treatment. Cultures treated with sepiolite and friction without the addition of 2-2′ bipyridyl and bacteria alone without sepiolite were used as a control. The mutant frequencies for these groups were determined as described elsewhere in this section.

### Assessing double-strand breaks with a plasmid system

To evaluate if sepiolite under friction treatment induces double-strand breaks in plasmid DNA, the strain *Escherichia coli* DH5α (fhuA2 lac(del)U169 phoA glnV44 Φ80’ lacZ(del)M15 gyrA96 recA1 relA1 endA1 thi-1 hsdR17) carrying the plasmid pET-19b (Novagen, Germany) was treated with sepiolite and sliding friction forces during one minute. Several samples were recovered from the plates and pooled to compensate viability losses due to friction. The recovery was done by washing the surface with 5 ml 0.9 % NaCl saline solution four times as described for mutagenesis experiments. The recovered pellets were washed with 1 ml of TE buffer and the OD_600_ adjusted to 1 for each type of sample. Plasmid DNA samples were extracted using a Qiagen mini plasmid extraction kit (Qiagen, Germany). Added sepiolite with or without friction and no sepiolite groups were used as a control group. Each experiment consisted of five replicates. The same amount of plasmid DNA per replicate was applied per well to an agarose gel that was stained with SYBR® Gold Nucleic Acid Gel Stain kit (Molecular Probes, USA). A NdeI (Promega, USA) digested aliquot of pET-19b was used as control of linear migration rate. The proportion of linear molecules of the plasmid were compared among groups using a densitometry analysis by ImageJ [55].

### RecA deficient strain construction

The *recA* null mutant was constructed following a previously described methodology [56] with the primers 5′-CAGAACATATTGACTATCCGGTATTACCCG-GCAT GACAGGAGT AAAAAT GGT-GT AGGCT GGAGCT GCTTC-3′ and 5-ATGCGACCCTTGTGTATCAAACAAGACGATTAAAAATCTTCGTTAGTTTCATGGGAAT-TAGCCATGGTCC-3′ (forward and reverse respectively) using the pKD3 plasmid as template. The mutant was checked by PCR amplification using the primers c1 5′-TTATACGCAAGGCGACAAGG-3′ and c2 5′-GATCTTCCGTCACAGGTAGG-3′ in combination with specific primers for upstream and downstream regions of *recA* gene: 5′-ATTGCAGACCTTGTGGCAAC-3′ and 5′-CGATCCAACAGGCGAGCATAT-3′ respectively. Additionally, the increased susceptibility to UV light and mitomycin C was tested phenotypically in comparison to the parental strain. The antibiotic resistance gene was eliminated using the pCP20 plasmid as described previously [56].

### SEM of *E. coli* treated with sepiolite

Approximately 2×10^9^ CFU of stationary phase *E. coli* MG1655 were treated with sepiolite and friction force was applied for one minute as described for the mutagenesis experiment. Circular agar blocks were taken from agar plates with a sterile cork borer (1 cm of diameter). Then, a thin surface layer was cut off, placed on a circular glass cover slip (1.5 cm of diameter) and incubated for 45 minutes at room temperature in a laminar flow cabinet to allow air drying of the samples. The cover glasses with dehydrated agar sections were mounted on aluminium stubs using double-sided adhesive tape and coated with gold in a sputter coater (SCD-040; Balzers, Union, Liechtenstein). The specimens were examined with a FEI Quanta 200 SEM (FEI Co., Hillsboro, OR) operating at an accelerating voltage of 15 kV under high vacuum mode at different magnifications. At least 5 sections from independent plates were observed to check physical penetration by the mineral. Some samples of sepiolite or asbestos (crocidotiles) alone were processed and observed in the same way.

### Long fibre-depleted sepiolite mutagenesis experiment

To assess the role of long fibre of sepiolite in mutagenesis, a sepiolite preparation depleted of fibres longer than 1 μm was obtained. A 100 ml sepiolite suspension (1 mg/ml) in distilled water was passed though Pall® Acrodisc® glass fibre syringe filters (Sigma, USA) several times. The resulting suspension was desiccated by evaporation at 70°C overnight. A non-filtered solution was used as a control. From the obtained powder, two suspensions were prepared to a final proportion of 0.1 mg/ml. These two solutions were used for a mutagenesis experiment plating in fosfomycin as indicated previously, using a friction time of two minutes.

### Mutant frequency estimation of asbestos treated cells

The procedure was carried out identically that the one described for sepiolite in this section. The time was set to two minutes and the same concentration that was used, 0.1 mg/ml. We used the crocidolite asbestos analytical standard (SPI Supplies, USA). The asbestos fibres were resuspended in distilled water, autoclaved and sonicated in bath during 10 minutes before use to render a homogeneous suspension.

### Statistical analysis

To compare experimental groups, Kruskal-Wallis test or One-way ANOVA test were performed. In case of significance, Bonferroni-corrected one-tailed Mann-Whitney U test or Tukey HSD Test were used respectively. P values less than or equal to 0.05, after correction if needed, were considered statistically significant. All tests were performed with the statistic software R v. 3.4.2 [57].

## Acknowledgements

ARR was supported by The Collaborative Research Centre (CRC) via project 973 “Priming and Memory of Organismic Responses to Stress”, project C5. J.B. was supported by Plan Nacional de I+D+i and Instituto de Salud Carlos III, Subdirección General de Redes y Centros de Investigación Cooperativa, Ministerio de Economía y Competitividad, Spanish Network for Research in Infectious Diseases (REIPI RD15/0012)-co-financed by European Development Regional Fund “A way to achieve Europe” ERDF and Fondo de Investigación Sanitaria Grant PI13/00063. We are grateful to Alejandro Couce, Olga Makarova and Jens Rolff for critical comments about the manuscript.

---

**Fig S1.**
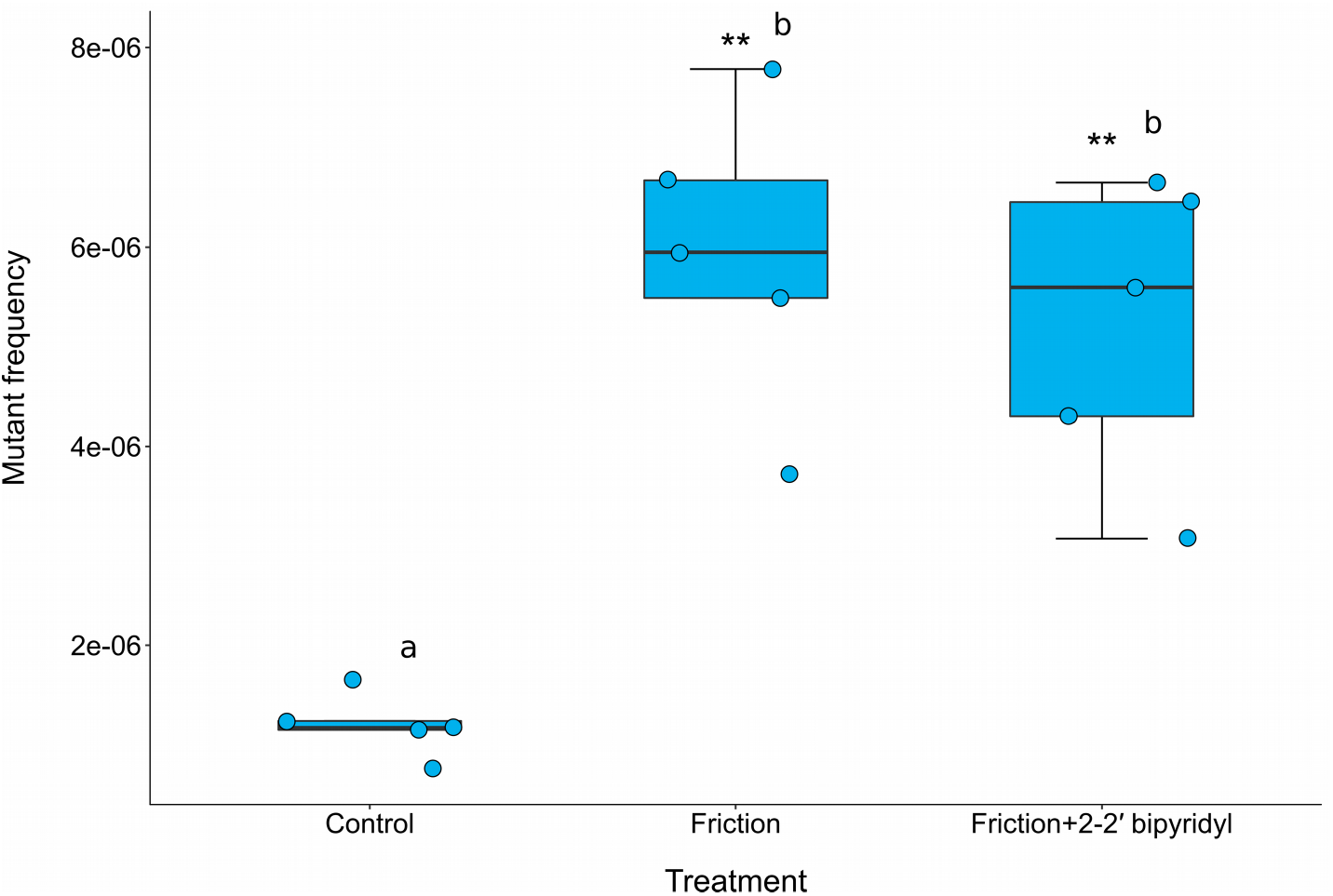
Addition of a chelating agent (2-2′ bipyridyl) does not significantly suppress or diminish the mutagenic effect of sepiolite. Box-plot of mutant frequency of *E. coli* MG1655 when added 2-2′ bipyridyl as chelating agent. Asterisks represents significant difference; Mann-Whitney *U*: P < 0.01. Equal letters represent no differences while different ones represent significant differences.

**Fig S2.**
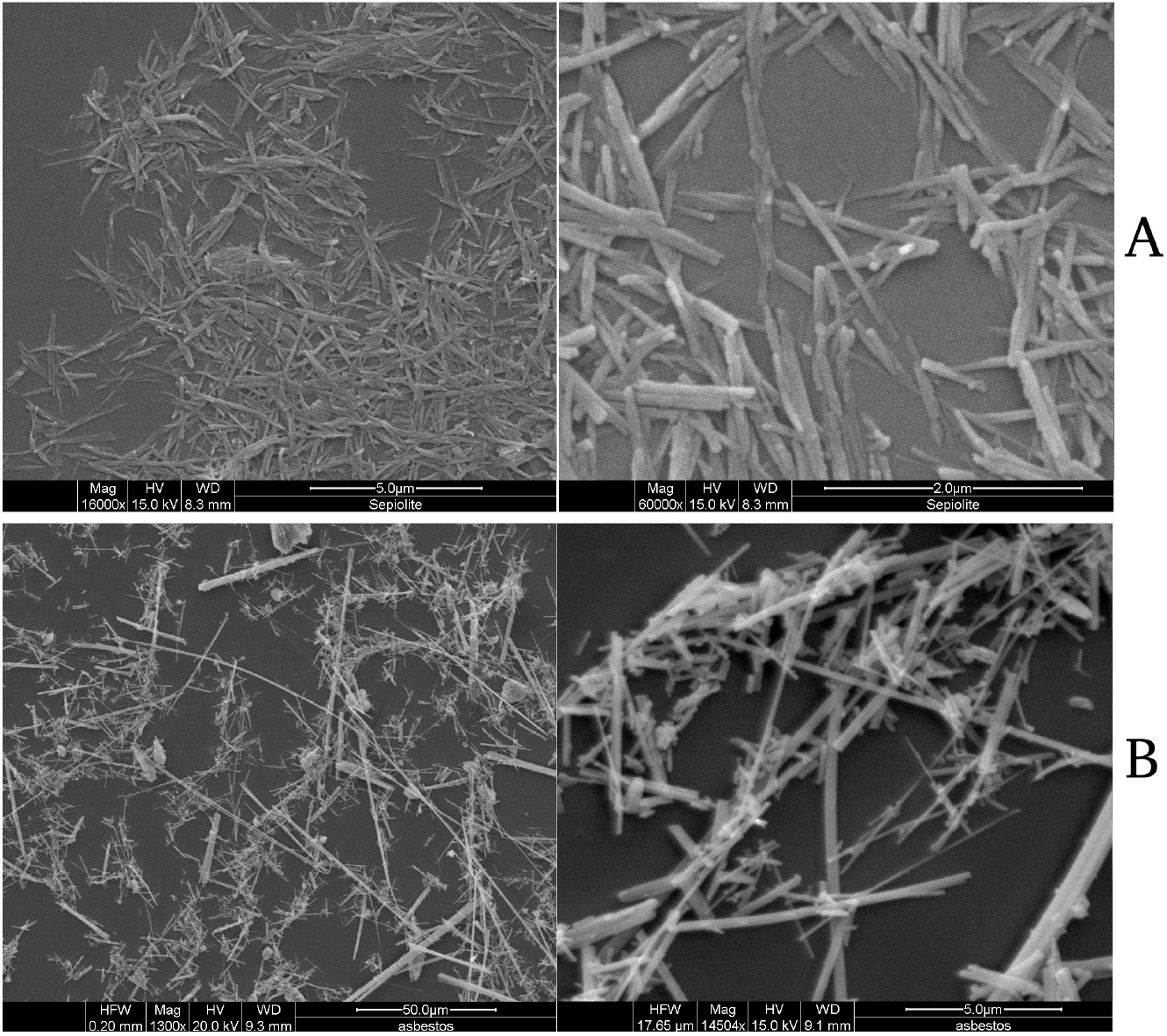
Visualisation of sepiolite and asbestos fibres under SEM. SEM examination of (A) sepiolite fibres and (B) asbestos. Fibres were observed at different magnifications as indicated in the pictures.

## References

1. Ouhida I, Pérez JF, Piedrafita J, Gasa J (2000) The effects of sepiolite in broiler chicken diets of high, medium and low viscosity. Productive performance and nutritive value. Anim Feed Sci Technol 85: 183–194.

2. Parisini P, Martelli G, Sardi L, Escribano F (1999) Protein and energy retention in pigs fed diets containing sepiolite. Anim Feed Sci Technol 79: 155–162. Available: http://www.animalfeedscience.com/article/S0377840199000085/fulltext. Accessed 18 June 2013.

3. Viseras C, Lopez-Galindo A (1999) Pharmaceutical applications of some spanish clays (sepiolite, palygorskite, bentonite): some preformulation studies. Appl Clay Sci 14: 69–82. Available: http://dx.doi.org/10.1016/S0169-1317(98)00050-7. Accessed 18 June 2013.

4. Rodríguez-Rojas A, Rodríguez-Beltrán J, Valverde JR, Blázquez J (2015) Can Clays in Livestock Feed Promote Antibiotic Resistance and Virulence in Pathogenic Bacteria? Antibiotics 4: 299–308. Available: http://www.mdpi.com/2079-6382/4/3/299. Accessed 16 July 2015.

5. Rodríguez-Beltrán J, Rodríguez-Rojas A, Yubero E, Blázquez J (2013) The Animal Food Supplement Sepiolite Promotes a Direct Horizontal Transfer of Antibiotic Resistance Plasmids between Bacterial Species. Antimicrob Agents Chemother 57: 2651–2653. Available: http://www.pubmedcentral.nih.gov/articlerender.fcgi?artid=3716135&tool=pmcentrez&rendertype=abstract. Accessed 18 June 2013.

6. Yoshida N, Sato M (2009) Plasmid uptake by bacteria: a comparison of methods and efficiencies. Appl Microbiol Biotechnol 83: 791–798. Available: http://www.ncbi.nlm.nih.gov/entrez/query.fcgi?cmd=retrieve&db=pubmed&dopt=citation&list_uids=19471921.

7. Dong L, Johnson DT (2005) Adsorption of acicular particles at liquid-fluid interfaces and the influence of the line tension. Langmuir 21: 3838–3849. Available: http://www.ncbi.nlm.nih.gov/entrez/query.fcgi?cmd=retrieve&db=pubmed&dopt=citation&list_uids=15835945. Accessed 18 June 2013.

8. Djordjevic SP, Stokes HW, Roy Chowdhury P (2013) Mobile elements, zoonotic pathogens and commensal bacteria: conduits for the delivery of resistance genes into humans, production animals and soil microbiota. Front Microbiol 4: 86. Available: http://www.pubmedcentral.nih.gov/articlerender.fcgi?artid=3639385&tool=pmcentrez&rendertype=abstract. Accessed 10 June 2013.

9. Wilharm G, Lepka D, Faber F, Hofmann J, Kerrinnes T, et al. (2010) A simple and rapid method of bacterial transformation. J Microbiol Methods 80: 215–216. Available: http://dx.doi.org/10.1016/j.mimet.2009.12.002. Accessed 18 June 2013.

10. Appel JD, Fasy TM, Kohtz DS, Kohtz JD, Johnson EM (1988) Asbestos fibers mediate transformation of monkey cells by exogenous plasmid DNA. Proc Natl Acad Sci U S A 85: 7670–7674. Available: http://www.pubmedcentral.nih.gov/articlerender.fcgi?artid=282254&tool=pmcentrez&rendertype=abstract. Accessed 1 July 2013.

11. Huang SXL, Jaurand M-C, Kamp DW, Whysner J, Hei TK (2011) Role of mutagenicity in asbestos fiber-induced carcinogenicity and other diseases. J Toxicol Environ Health B Crit Rev 14: 179–245. Available: http://www.pubmedcentral.nih.gov/articlerender.fcgi?artid=3118525&tool=pmcentrez&rendertype=abstract. Accessed 6 April 2016.

12. Daniel FB (1983) In vitro assessment of asbestos genotoxicity. Environ Health Perspect 53: 163–167. Available: http://www.ncbi.nlm.nih.gov/pubmed/6363052. Accessed 2 November 2016.

13. Fowler P, Homan A, Atkins D, Whitwell J, Lloyd M, et al. (2016) The utility of the in vitro micronucleus test for evaluating the genotoxicity of natural and manmade nano-scale fibres. Mutat Res Toxicol Environ Mutagen 809: 33–42. Available: http://www.ncbi.nlm.nih.gov/pubmed/27692297. Accessed 6 January 2017.

14. Jaurand MC (1997) Mechanisms of fiber-induced genotoxicity. Environ Health Perspect 105 Suppl: 1073–1084. Available: http://www.pubmedcentral.nih.gov/articlerender.fcgi?artid=1470173&tool=pmcentrez&rendertype=abstract. Accessed 15 April 2016.

15. Berendonk TU, Manaia CM, Merlin C, Fatta-Kassinos D, Cytryn E, et al. (2015) Tackling antibiotic resistance: the environmental framework. Nat Rev Microbiol 13: 310–317. Available: http://dx.doi.org/10.1038/nrmicro3439. Accessed 22 November 2015.

16. Blázquez J, Couce A, Rodríguez-Beltrán J, Rodríguez-Rojas A, Blazquez J, et al. (2012) Antimicrobials as promoters of genetic variation. Curr Opin Microbiol 15: 561–569. Available: http://www.ncbi.nlm.nih.gov/pubmed/22890188. Accessed 24 May 2013.

17. Zipperer A, Konnerth MC, Laux C, Berscheid A, Janek D, et al. (2016) Human commensals producing a novel antibiotic impair pathogen colonization. Nature 535: 511–516. Available: http://www.nature.com/doifinder/10.1038/nature18634. Accessed 6 January 2017.

18. Williams LB, Metge DW, Eberl DD, Harvey RW, Turner AG, et al. (2011) What makes a natural clay antibacterial? Environ Sci Technol 45: 3768–3773. Available: /pmc/articles/PMC3126108/?report=abstract. Accessed 26 June 2015.

19. Yoshida N, Ide K (2008) Plasmid DNA is released from nanosized acicular material surface by low molecular weight oligonucleotides: exogenous plasmid acquisition mechanism for penetration intermediates based on the Yoshida effect. Appl Microbiol Biotechnol 80: 813–821. Available: http://www.ncbi.nlm.nih.gov/pubmed/18704395. Accessed 18 June 2013.

20. Lieber MR (2010) The mechanism of double-strand DNA break repair by the nonhomologous DNA end-joining pathway. Annu Rev Biochem 79: 181–211. Available: http://www.ncbi.nlm.nih.gov/pubmed/20192759. Accessed 11 July 2014.

21. Sambrook J, Russell DW (2001) Molecular cloning: a laboratory manual. 3rd ed. Cold Spring Harbor [New York]: Laboratory Press.

22. Persky NS, Lovett ST (2008) Mechanisms of Recombination: Lessons from E. coli. Crit Rev Biochem Mol Biol 43: 347–370. doi:10.1080/10409230802485358.

23. Akerlund T, Nordström K, Bernander R (1995) Analysis of cell size and DNA content in exponentially growing and stationary-phase batch cultures of Escherichia coli. J Bacteriol 177: 6791–6797. Available: http://www.ncbi.nlm.nih.gov/pubmed/7592469. Accessed 6 January 2017.

24. Ponder RG, Fonville NC, Rosenberg SM (2005) A switch from high-fidelity to error-prone DNA double-strand break repair underlies stress-induced mutation. Mol Cell 19: 791–804. Available: http://www.ncbi.nlm.nih.gov/pubmed/16168374. Accessed 3 April 2016.

25. Shee C, Gibson JL, Darrow MC, Gonzalez C, Rosenberg SM (2011) Impact of a stressinducible switch to mutagenic repair of DNA breaks on mutation in Escherichia coli. Proc Natl Acad Sci U S A 108: 13659–13664. Available: http://www.ncbi.nlm.nih.gov/pubmed/21808005. Accessed 14 November 2016.

26. Boles BR, Thoendel M, Singh PK (2004) Self-generated diversity produces “insurance effects” in biofilm communities. Proc Natl Acad Sci U S A 101: 16630–16635. Available: http://www.ncbi.nlm.nih.gov/entrez/query.fcgi?cmd=retrieve&db=pubmed&dopt=citation&list_uids=15546998.

27. Rosenberg SM, Shee C, Frisch RL, Hastings PJ (2012) Stress-induced mutation via DNA breaks in Escherichia coli: a molecular mechanism with implications for evolution and medicine. Bioessays 34: 885–892. Available: http://www.ncbi.nlm.nih.gov/pubmed/22911060. Accessed 6 January 2017.

28. Dorman CJ (2013) Genome architecture and global gene regulation in bacteria: making progress towards a unified model? Nat Rev Microbiol 11: 349–355. Available: http://dx.doi.org/10.1038/nrmicro3007. Accessed 6 April 2016.

29. Wolf SG, Frenkiel D, Arad T, Finkel SE, Kolter R, et al. (1999) DNA protection by stress-induced biocrystallization. Nature 400: 83–85. Available: http://www.ncbi.nlm.nih.gov/pubmed/10403254. Accessed 12 April 2016.

30. Pelletier J, Halvorsen K, Ha B-Y, Paparcone R, Sandler SJ, et al. (2012) Physical manipulation of the Escherichia coli chromosome reveals its soft nature. Proc Natl Acad Sci U S A 109: E2649–56. Available: http://www.ncbi.nlm.nih.gov/pubmed/22984156. Accessed 5 January 2017.

31. Eastmond DA, Hartwig A, Anderson D, Anwar WA, Cimino MC, et al. (2009) Mutagenicity testing for chemical risk assessment: update of the WHO/IPCS Harmonized Scheme. Mutagenesis 24: 341–349. Available: http://www.ncbi.nlm.nih.gov/pubmed/19535363. Accessed 28 October 2016.

32. Yoshida N, Naka T, Ohta K (2004) Mutagenesis of bacteria by fibrous or clay minerals. J Biol Sci 4: 532–526. Available: http://www.scialert.net/abstract/?doi=jbs.2004.532.536. Accessed 15 November 2016.

33. Barlow CA, Lievense L, Gross S, Ronk CJ, Paustenbach DJ (2013) The role of genotoxicity in asbestos-induced mesothelioma: an explanation for the differences in carcinogenic potential among fiber types. Inhal Toxicol 25: 553–567. Available: http://www.ncbi.nlm.nih.gov/pubmed/23905972. Accessed 15 April 2016.

34. Miserocchi G, Sancini G, Mantegazza F, Chiappino G (2008) Translocation pathways for inhaled asbestos fibers. Environ Health 7: 4. Available: http://www.ncbi.nlm.nih.gov/pubmed/18218073. Accessed 2 November 2016.

35. Rodriguez-Beltran J, Elabed H, Gaddour K, Blazquez J, Rodriguez-Rojas A (2012) Simple DNA transformation in Pseudomonas based on the Yoshida effect. J Microbiol Methods 89: 95–98. Available: http://www.ncbi.nlm.nih.gov/entrez/query.fcgi?cmd=retrieve&db=pubmed&dopt=citation&list_uids=22405834.

36. Tan H, Fu L, Seno M (2010) Optimization of bacterial plasmid transformation using nanomaterials based on the yoshida effect. Int J Mol Sci 11: 4961–4972. Available: http://www.pubmedcentral.nih.gov/articlerender.fcgi?artid=3100829&tool=pmcentrez&rendertype=abstract. Accessed 1 July 2013.

37. Mitsudome Y, Takahama M, Hirose J, Yoshida N (2014) The use of nano-sized acicular material, sliding friction, and antisense DNA oligonucleotides to silence bacterial genes. AMB Express 4: 70. Available: http://www.pubmedcentral.nih.gov/articlerender.fcgi?artid=4230895&tool=pmcentrez&rendertype=abstract. Accessed 6 April 2016.

38. Toyokuni S (2013) Genotoxicity and carcinogenicity risk of carbon nanotubes. Adv Drug Deliv Rev 65: 2098–2110. Available: http://www.ncbi.nlm.nih.gov/pubmed/23751780. Accessed 15 April 2016.

39. Chernova T, Murphy FA, Galavotti S, Sun X-M, Powley IR, et al. (2017) Long-Fiber Carbon Nanotubes Replicate Asbestos-Induced Mesothelioma with Disruption of the Tumor Suppressor Gene Cdkn2a (Ink4a/Arf). Curr Biol 27: 3302–3314.e6. Available: http://www.ncbi.nlm.nih.gov/pubmed/29112861. Accessed 8 January 2018.

40. Rojas-Chapana J, Troszczynska J, Firkowska I, Morsczeck C, Giersig M (2005) Multi-walled carbon nanotubes for plasmid delivery into Escherichia coli cells. Lab Chip 5: 536–539. Available: http://www.ncbi.nlm.nih.gov/pubmed/15856091. Accessed 9 March 2016.

41. Yoshida N, Kodama K, Nakata K, Yamashita M, Miwa T (2002) Escherichia coli cells penetrated by chrysotile fibers are transformed to antibiotic resistance by incorporation of exogenous plasmid DNA. Appl Microbiol Biotechnol 60: 461–468. Available: http://www.ncbi.nlm.nih.gov/pubmed/12466888. Accessed 1 July 2013.

42. Rodríguez-Beltrán J, Elabed H, Gaddour K, Blázquez J, Rodríguez-Rojas A (2012) Simple DNA transformation in Pseudomonas based on the Yoshida effect. J Microbiol Methods 89: 95–98. Available: http://www.ncbi.nlm.nih.gov/pubmed/22405834. Accessed 18 June 2013.

43. Terry KL, Karageorgi S, Shvetsov YB, Merritt MA, Lurie G, et al. (2013) Genital powder use and risk of ovarian cancer: a pooled analysis of 8,525 cases and 9,859 controls. Cancer Prev Res (Phila) 6: 811–821. Available: http://www.ncbi.nlm.nih.gov/pubmed/23761272. Accessed 18 November 2016.

44. Muscat JE, Huncharek MS (2008) Perineal talc use and ovarian cancer: a critical review. Eur J Cancer Prev 17: 139–146. Available: http://www.ncbi.nlm.nih.gov/pubmed/18287871. Accessed 18 November 2016.

45. Anderson EL, Sheehan PJ, Kalmes RM, Griffin JR (2016) Assessment of Health Risk from Historical Use of Cosmetic Talcum Powder. Risk Anal. Available: http://doi.wiley.com/10.1111/risa.12664. Accessed 18 November 2016.

46. Wu W-N, McKown LA (2004) In Vitro Drug Metabolite Profiling Using Hepatic S9 and Human Liver Microsomes. Optimization in Drug Discovery. Totowa, NJ: Humana Press. pp. 163–184. Available: http://link.springer.com/10.1385/1-59259-800-5:163. Accessed 18 November 2016.

47. Yong E (2013) Gut microbes keep species apart. Nature. Available: http://www.nature.com/news/gut-microbes-keep-species-apart-1.13408?WT.ec_id=NEWS-20130723. Accessed 5 April 2016.

48. Wings O (2007) A review of gastrolith function with implications for fossil vertebrates and a revised classification - Acta Palaeontologica Polonica. Acta Palaeontol Pol 52: 1–16. Available: https://www.app.pan.pl/article/item/app52-001.html. Accessed 5 April 2016.

49. Hansson GC (2012) Role of mucus layers in gut infection and inflammation. Curr Opin Microbiol 15: 57–62. Available: /pmc/articles/PMC3716454/?report=abstract. Accessed 29 October 2015.

50. Barrett JC, Lamb PW, Wiseman RW (1989) Multiple mechanisms for the carcinogenic effects of asbestos and other mineral fibers. Environ Health Perspect 81: 81–89. Available: http://www.ncbi.nlm.nih.gov/pubmed/2667990. Accessed 3 November 2016.

51. Foley-Comer AJ, Herrick SE, Al-Mishlab T, Prêle CM, Laurent GJ, et al. (2002) Evidence for incorporation of free-floating mesothelial cells as a mechanism of serosal healing. J Cell Sci 115: 1383–1389. Available: http://www.ncbi.nlm.nih.gov/pubmed/11896186. Accessed 18 November 2016.

52. Mendes GP, Vieira PS, Lanceros-Méndez S, Kluskens LD, Mota M (2015) Transformation of Escherichia coli JM109 using pUC19 by the Yoshida effect. J Microbiol Methods 115: 1–5. Available: http://linkinghub.elsevier.com/retrieve/pii/S0167701215001517. Accessed 3 November 2016.

53. Hall BM, Ma C-X, Liang P, Singh KK (2009) Fluctuation analysis CalculatOR: a web tool for the determination of mutation rate using Luria-Delbruck fluctuation analysis. Bioinformatics 25: 1564–1565. Available: http://www.pubmedcentral.nih.gov/articlerender.fcgi?artid=2687991&tool=pmcentrez&rendertype=abstract. Accessed 1 May 2014.

54. Kaes C, Katz A, Hosseini MW (2000) Bipyridine: The Most Widely Used Ligand. A Review of Molecules Comprising at Least Two 2,2‘-Bipyridine Units. Chem Rev 100: 3553–3590. Available: http://dx.doi.org/10.1021/cr990376z. Accessed 6 April 2016.

55. Schneider CA, Rasband WS, Eliceiri KW (2012) NIH Image to ImageJ: 25 years of image analysis. Nat Methods 9: 671–675. Available: http://dx.doi.org/10.1038/nmeth.2089. Accessed 9 July 2014.

56. Datsenko KA, Wanner BL (2000) One-step inactivation of chromosomal genes in Escherichia coli K-12 using PCR products. Proc Natl Acad Sci U S A 97: 6640–6645. Available: http://www.ncbi.nlm.nih.gov/entrez/query.fcgi?cmd=retrieve&db=pubmed&dopt=citation&list_uids=10829079.

57. R Development Core Team (n.d.) R Development Core Team (2013). R: A language and environment for statistical computing. R Foundation for Statistical Computing, Vienna, Austria. I ISBN 3-900051-07-0, URL http://www.R-project.org.

